# Knockdown of *PHYTOENE DESATURASE* in the field dodder, *Cuscuta campestris*, by Virus-Induced Gene Silencing

**DOI:** 10.1101/2021.04.07.438779

**Authors:** Steven Dyer, Ryan Weir, Panagiotis Manesiotis, Johnathan J. Dalzell

## Abstract

*Cuscuta campestris* is a globally distributed obligate holoparasitic plant, and economically important crop pest. There is an urgent need for safe and effective new herbicides to control *Cuscuta* spp. PHYTOENE DESATURASE (PDS) is a biosynthetic enzyme within the carotenoid synthesis pathway, which is a target for several commercially available herbicides. The low transpiration rate of *C. campestris* results in sub-optimal translocation of PDS-targeting herbicides throughout the parasite, and resistance to these herbicides, and others, should be anticipated. Here we demonstrate that RNA interference (RNAi) can effectively reduce the expression of *PDS* in *C. campestris*. Virus Induced Gene Silencing (VIGS) is capable of inducing *PDS* knockdown in *C. campestris*, when Tobacco Rattle Virus (TRV) is used to deliver a *PDS*-specific sequence through the host plant *Arabidopsis thaliana*. This leads to a reduction in the accumulation of beta carotene, which is synthesised from phytoene, and significantly reduced growth of *C. campestris*. We hypothesise that secondary amplification and spread of *PDS* double-stranded RNA within *C. campestris* may circumvent the translocation limitations of other xylem and phloem-spread PDS-specific herbicides. These data demonstrate for the first time that VIGS can be used for reverse genetics interrogation of the *C. campestris* genome.

## Introduction

The field dodder, *Cuscuta campestris*, is a globally distributed flowering plant, and obligate holoparasite of crops. Seeds form within indehiscent fruits, which can be dispersed by endozoochory (Costea et al., 2016), and may remain viable for decades through physical dormancy (Gaertner, 1950; Goss, 1924). Endozoochoric dispersal has the potential to cause sporadic and unpredictable infestation events, and dodder infection can also spread vegatatively within and between neighbouring plants. Dodder seedlings locate host plants through phototropism (Johnson et al., 2016), olfaction (Runyon et al., 2006) and mechanosensation (Kastier et al., 2018), which is further aided by the natural circumnutation of dodder shoot growth (Johnson et al., 2016). Once a host plant stem has been located, thigmotropic growth promotes coiling of the dodder shoot around the host stem. The perception of mechanical stimulation and far-red light, which is indicative of canopy shading, is sufficient to trigger shoot adhesion, and haustorium organogenesis (Shimizu and Aoki, 2019). The haustorium facilitates attachment to, and subsequent invasion of the host. Haustorium cells will penetrate the host stem, and fuse with host phloem and xylem cells, allowing the withdrawal of water and other metabolic resources. Dodder parasitism has a significant negative impact on global crop productivity, especially for small-hold subsistence farmers (Masanga et al., 2020). In Kenya, the Kikuyu refer to dodder as the ‘poverty plant’.

*C. campestris* forms tangled vines within and between host plants, which makes mechanical weeding extremely difficult. *C. campestris* is also naturally resistant to several commonly used herbicides (Nadler-Hassar et al., 2009). However, it is possible for *C. campestris* to acquire resistance following infection of a glufosinate-resistant host crop. Several herbicides target PHYTOENE DESATURASE (PDS), which is an enzyme involved in the synthesis of carotenoids. Although some of these herbicides are effective for dodder control, there are inherent limitations due to the translocation characteristics of the respective compounds, corresponding to the low rate of transpiration in *C. campestris* (Weinberg et al., 2003). This underlines the complexity and challenges of dodder control and highlights the need to develop effective new approaches. The genome sequence of two *C. campestris* populations have recently been published (Yang et al., 2019; Vogel et al., 2018), providing new opportunities for the rational design of targeted herbicides.

Phytoene is a precursor in the synthesis of the orange carotenoid pigment, β-carotene, which is itself a precursor for the synthesis of several hormones, including strigolactones (reviewed by Nisar et al., 2015). Genes involved in strigolactone signalling are upregulated during pre-haustorium development in *Cuscuta* spp. (Ranjan et al., 2014), which suggests that interfering with PDS function may have knock-on effects relevant to strigolactone synthesis and the induction of haustorium organogenesis. Strigolactones can also promote or inhibit shoot branching, depending on auxin signalling events (Shinohara et al., 2013). Abscisic acid (ABA) transport is altered during the pre-haustorium development stage, and β-carotene is a precursor for ABA synthesis (Nisar et al., 2015). Carotenoid pigments that are synthesised from phytoene are also important for photoprotection; the metabolic quenching of reactive oxygen species that are generating from light processing (Snyder et al., 2005; Bungard et al., 1999). Knowing that PDS is a viable target for dodder control, we aimed to establish the efficacy of *PDS* gene knockdown through RNA interference (RNAi), as a potential approach for dodder management, and to validate the potential of RNAi for the study of gene function in *C. campestris*.

RNAi is a biochemical pathway that directs gene knockdown via target-specific small non-coding RNAs (Hamilton and Baulcombe, 1999; Napoli et al., 1990). Host-induced gene silencing (HIGS) is capable of inducing gene knockdown in *Cuscuta pentagona* (Alakonya et al., 2012), and the field dodder, *C. campestris* (Jhu et al., 2021). Virus-Induced Gene Silencing (VIGS) represents a rapid, high-throughput approach to triggering RNAi, which exploits the natural defence response of plants to viral RNA sequences (Burch-Smith et al., 2006). Here we aimed to determine if VIGS could be used to silence the *PDS* gene in *C. campestris* during parasitism of the model host plant *Arabidopsis thaliana*. The development of a rapid and reliable VIGS protocol for the study of *C. campestris* gene function would greatly expedite the study of *C. campestris* biology and host-parasite interactions.

## Materials and methods

### Plant culture and maintenance

Seeds of *A. thaliana* Col-0 were surface sterilised in 1% sodium hypochlorite for 15 minutes and subsequently rinsed 5 times in ddH_2_O before being spread on 1 x MS media (1% agar, 4.3 g/L Murashige and skoog basal medium, 0.5 g/L MES, pH 5.7) in a 0.1% sterile agar suspension. Plates were stratified at 4° C in darkness for 48 h, then transferred to a walk-in plant growth room with full spectrum LED lights under long day conditions (16 h day / 8 h night) at a constant temperature of 23°C. Germinated seedlings with the first two true leaves were transferred to sterile Jack’s Magic All Purpose Compost (Westland). Plastic lids were placed over trays for 24 h to maintain humidity. As soon as *A. thaliana* plants had begun to bolt, *C. campestris* seeds were scarified and germinated.

Scarification of *C. campestris* seeds was achieved through incubation in 3M H_2_SO_4_ for 1 h followed by surface sterilisation in 1% sodium hypochlorite for two minutes, and five washes in ddH_2_O. *C. campestris* seeds were then placed on sterile filter paper, moistened with ddH_2_0, inside a standard Petri dish, and transferred to a 30°C incubator in darkness. Under these conditions, up to 100% of seeds would germinate after 72 h, which allowed for sufficient growth of *A. thaliana* host plants to facilitate *C. campestris* attachment (Figure 1). *C. campestris* seedlings were introduced to *A. thaliana* host plants individually and infection was facilitated by supplementing with far-red light (SolarSystem Far Red | 100 Watts, California Lightworks) for 72 h. This was sufficient to induce thigmotropic attachment, haustorium organogenesis and invasion. Once the far red light was removed, *C. campestris* was unable to form additional haustorial connections due to the altered light profile. All data have been generated using *C. campestris* seedlings that were cultured for at least two generations on *A. thaliana* (col-0).

**Figure 1.**
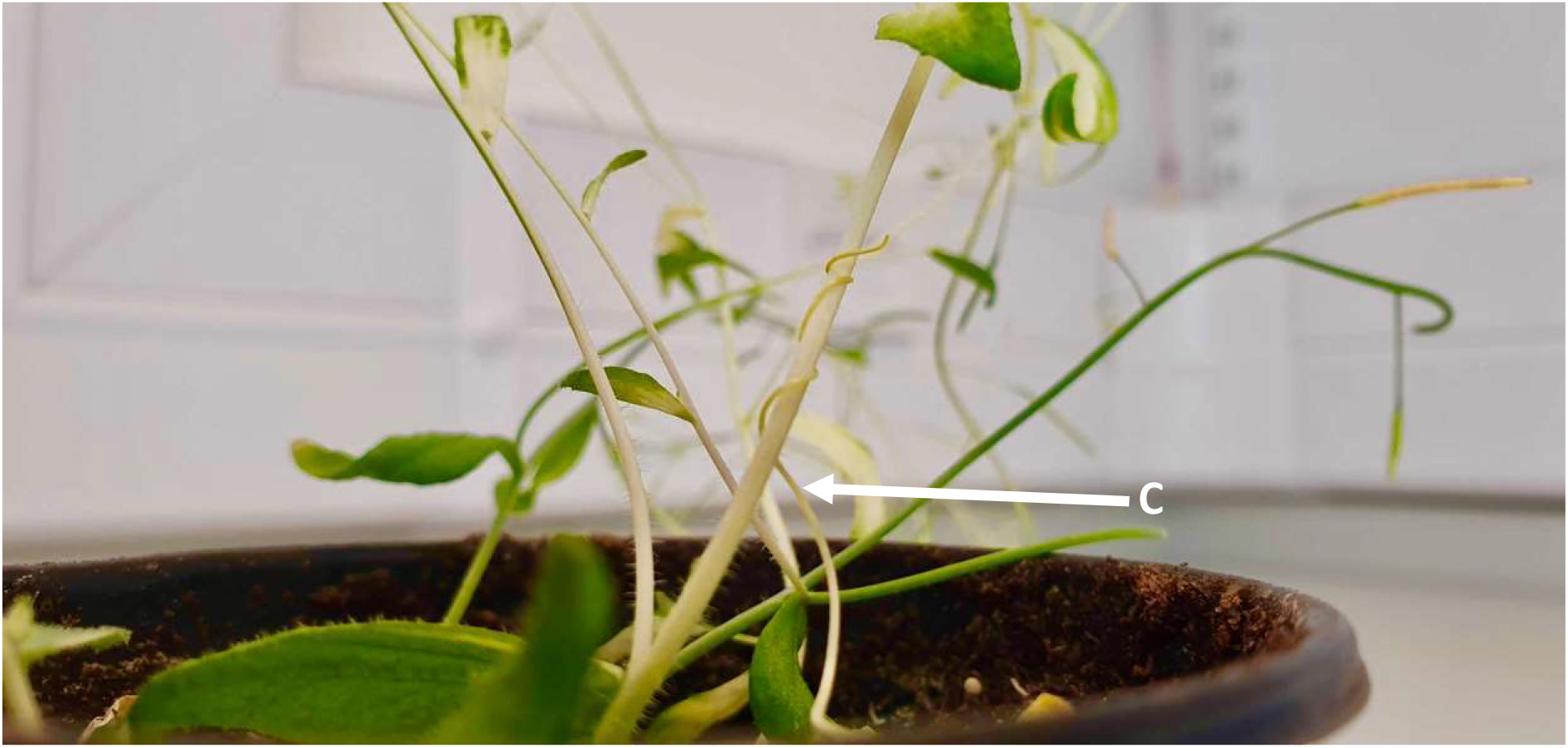
*C. campestris* coiling around a host plant. Unlike VIGS in tomato, silencing of the *PDS* gene in *A. thaliana* causes a pronounced photobleaching effect within the shoot and therefore site of parasite attachment. (C) Denotes the seedling of *C. campestris*.

### VIGS Plasmid design

Opting for a co-silencing system as previously described (Orzaez et al., 2009; Stratmann and Hind, 2011), tobacco rattle virus vectors were designed and transformed into *Agrobacterium tumefaciens* strain GV3101 as in Cox et al. (2019). A control construct, specific to a neuropeptide gene in the nematode *Meloidogyne incognita* (*Miflp16*), was also constructed. The 200 bp region for this construct shares no significant sequence similarity with either *A. thaliana* or *C. campestris* cDNA sequences. Sequence alignments for each of these sequences indicated less than 50% overall similarity between reporter, target and control and no more than 5 bp contiguous sequences across the entire target sequence (Table 1).

**Table 1.**
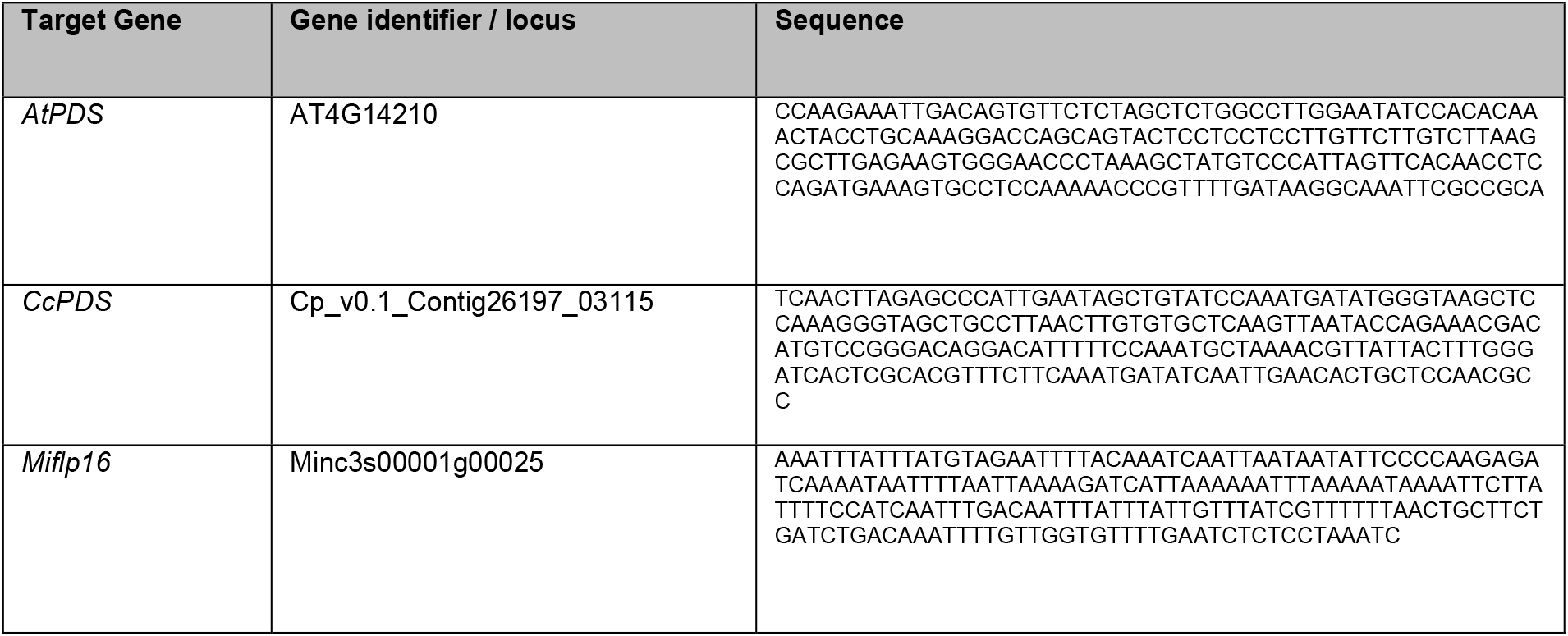
Insert sequences for pTRV2 VIGS vectors to silence *PDS* in *C. campestris*.

### Preparation of *A. tumefaciens* for plant inoculation

Frozen aliquots of *A. tumefaciens* containing either the pTRV1, pTRV2_*CcPDS* or pTRV2_*Miflp16* inserts were separately inoculated into 5 mL LB broth (Sigma) with 50 µg/mL gentamycin and 50 µg/mL kanamycin, three days prior to plant inoculation. Cultures were transferred to an orbital shaker, and maintained at 28°C for 48 h at 250 rpm. Subsequently, each 5 mL culture was diluted to 45 mL with LB broth supplemented with 10 mM morpholineethanesulfonic acid (MES), pH 5.7 and 20 µM acetosyringone (Sigma). Cultures were returned to the 28°C orbital shaker at 250 rpm overnight. On the day of plant inoculation, each culture was pelleted by centrifugation at 4000 x g for 10 minutes and resuspended to an OD_600_ of 1.5 in inoculation buffer (10 mM MES, 200 µm acetosyringone, 10 mM MgCl_2_) and incubated at room temperature in darkness for 3 hours.

### Plant inoculation and parasite infections

*A. thaliana* host plants were ready for inoculation approximately 2 weeks after the 4-6 leaf stage. Immediately prior to inoculation, the pTRV1 and pTRV2 cultures were mixed in a 1:1 ratio to give 10 mL of culture. A 21 G needle was used to pierce two leaves per plant. 1 mL of the premixed culture was drawn up using a needle-less syringe, and the aperture of the syringe was placed around the punctured hole on the abaxial surface of the leaf. The wound was sealed on the adaxial surface using a finger and gentle pressure was applied to the plunger until visual darkening of the leaf was observed, which confirmed successful infiltration into the leaf mesoderm. Generally less than 100 µl was sufficient per plant.

After inoculation, plants were returned to growth chambers and covered with a foil lined plastic lid for 24 h. The lids were removed after 24 h. Photobleaching of the leaves and stem occurred around two weeks following inoculation, at which time *C. campestris* seedlings were first introduced. *C. campestris* seeds were scarified 11 days after inoculation of *A. thaliana* host plants. *C. campestris* vines were sampled from experimental and control plants at 7, 14 and 21 days post attachment (dpa). Fresh weight and length measurements of *C. campestris* were measured, alongside target and non-target mRNA abundance. Tissue β-carotene abundance was also quantified.

### RT-qPCR analysis of gene knockdown

For *C. campestris* vines sampled at days 14 and 21, tissue was sampled at two distinct sites; the most proximal 6 cm immediately adjacent to the haustorium and the most distal 6 cm of the growing shoot, where presumptive haustoria would form. For samples at day 7, whole *C. campestris* vines were sampled as the length of most specimens was less than 12 cm. Freshly excised tissue was immediately frozen in liquid nitrogen and homogenised using the QIAGEN Tissue Lyser LT with a 5 mm stainless steel bead. RNA extraction and cDNA synthesis were performed as in Cox et al. (2019). cDNA was diluted 10 fold to 200 µL and 2 µl of cDNA was used for each qRT-PCR reaction in a total reaction volume of 12 µl, including 666 nM of each primer and 1x SensiFAST SYBR No-ROX mix (BIOLINE). Primers were designed to anneal outside the 200 bp VIGS target sequence (Table. 2). Technical PCR reactions were performed in triplicate for each target using a Rotorgene Q thermal cycler with the following cycle conditions: 95 °C × 10 min, 45 × (95 °C × 20 s, 60 °C × 20 s, 72 °C × 25 s) 72 °C × 10 min. PCR efficiencies of each amplicon, and the corresponding cT value were calculated using the Rotorgene Q software. Relative quantification of each target amplicon was obtained by an augmented comparative Ct method, relative to the ribosomal 18s housekeeper gene, as in Alakonya et al. (2012). Ratio-changes in transcript abundance were calculated relative to *C. campestris* tissue from non-inoculated control plants. Comparisons between *PDS* treated and *Miflp16* control treated plants was conducted by unpaired T-test (Graphpad Prism version 8).

**Table 2.**
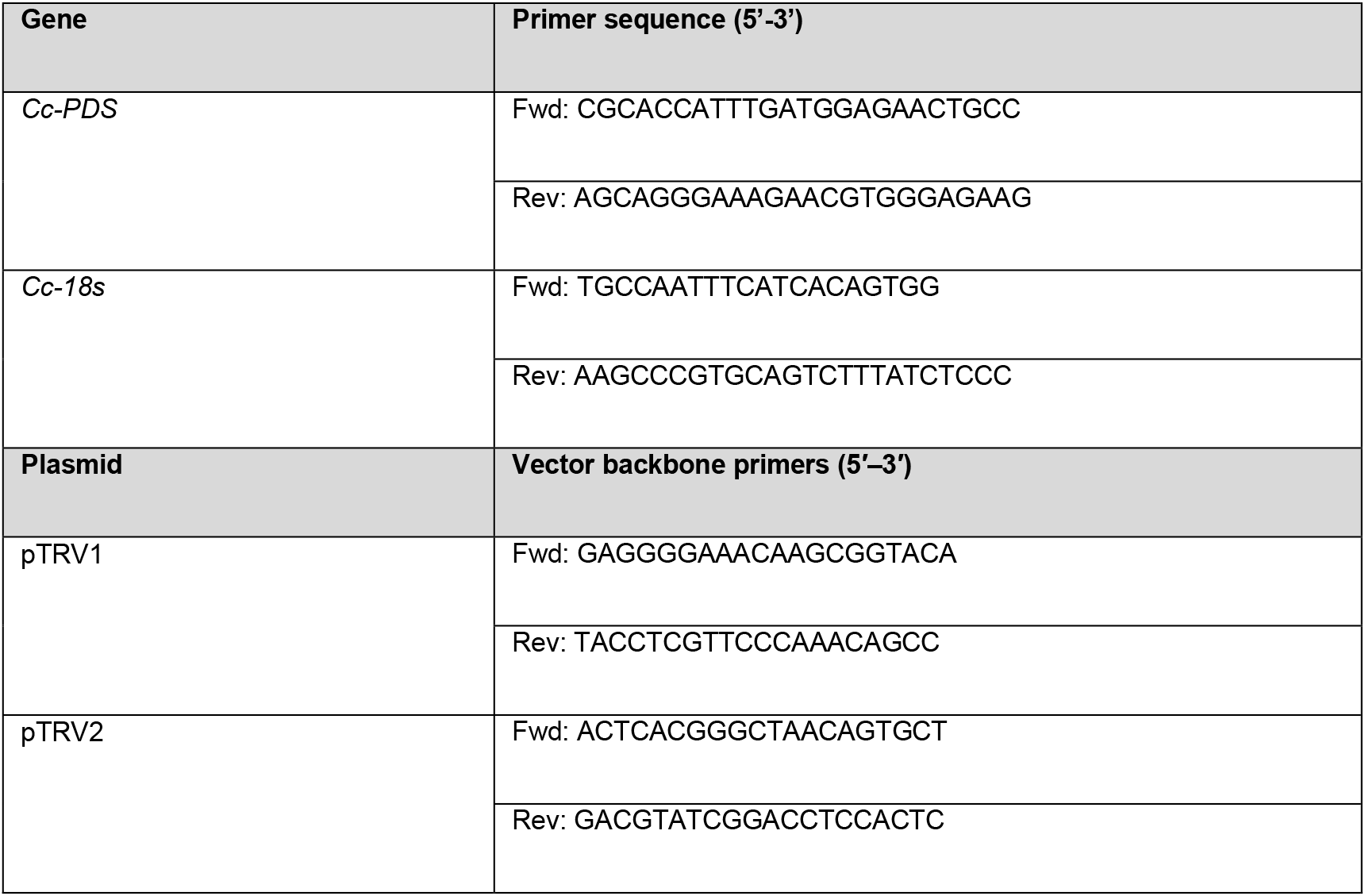
qRT-PCR primer sequences for *C. campestris*.

### Analysis of *C. campestris* growth

*C. campestris* was excised from the host *A. thaliana* after 7, 14 and 21 dpa. Fresh weight and length of each *C. campestris* plant was recorded. Length measurements of each *C. campestris* plant represents the length of the main stem, and all branched stems combined. Treatment groups were compared by one-way ANOVA and Tukey’s HSD post hoc test (Graphpad Prism version 8).

### Quantification of β-carotene

All solvents for β-carotene extraction (HPLC grade), β-carotene standards, and solvents for UPLC-MS analysis were from Sigma Aldrich (Gillingham, UK). Tissue samples were frozen in liquid nitrogen and ground to a fine powder using a pestle and mortar under liquid nitrogen and 100 mg of homogenised tissue was added to a 2 mL microcentrifuge tube. Pigments were isolated from tissue using a methodology modified from Gupta et al. (2015). 1.5 mL of chloroform:dichloromethane (2:1 v/v) was added to the homogenate and the suspension was incubated at 4 °C for 20 minutes with regular mixing by vortex every 5 minutes. Following incubation, 500 µL of 1 M sodium chloride was added and mixed by inversion. Phases were separated by centrifugation at 5,000 x g for 10 minutes and the organic phase was collected. The aqueous phase was re-extracted with 0.75 mL of chloroform:dichloromethane (2:1, v/v), centrifuged again and the organic phase was collected. Organic phases were pooled and subsequently dried by centrifugal evaporation at 30°C for 45 minutes. The pellet was redissolved in 10 mL chloroform:methanol (1:9 v/v). The samples were finally analysed on an Agilent 1290 UPLC system coupled to a photodiode array detector and a G6125B MSD, using a Poroshell 120 C18 column (50 x 2.1 mm, 2.7 μm). The mobile phase consisted of methanol with 0.1% formic acid with a flow rate of 0.4 mL min^-1^. Samples were ionised by API-ES in positive polarity and selected ion monitoring mode (SIM) set at m/z=536.40 for β-carotene. Quantification of β-carotene was conducted using a calibration curve previously prepared using β-carotene standards.

## Results

### Knockdown of *PHYTOENE DESATURDASE* by Virus-Induced Gene Silencing

At 7 dpa, whole vines were sampled and target transcript abundance was reduced by 71.80 ± 6.60 % relative to *flp16-*treated plants (Figure 2). At 14 and 21 dpa the vines were sampled in two discrete sections. The most proximal 6 cm to the site of attachment and the most distal 6 cm including the apical tip. We observed differences in target transcript abundance across the length of the vine. At 14 dpa, the proximal site there was a reduction of 51.60 % ± 8.85% whereas we observed no statistically significant knockdown at the distal site. Likewise, at 21 dpa there was no statistically significant *PDS* gene knockdown in the distal sections of *C. campestris* vines, however *PDS* transcript abundance was reduced by 32.17 ± 4.84 % in proximal tissues.

**Figure 2.**
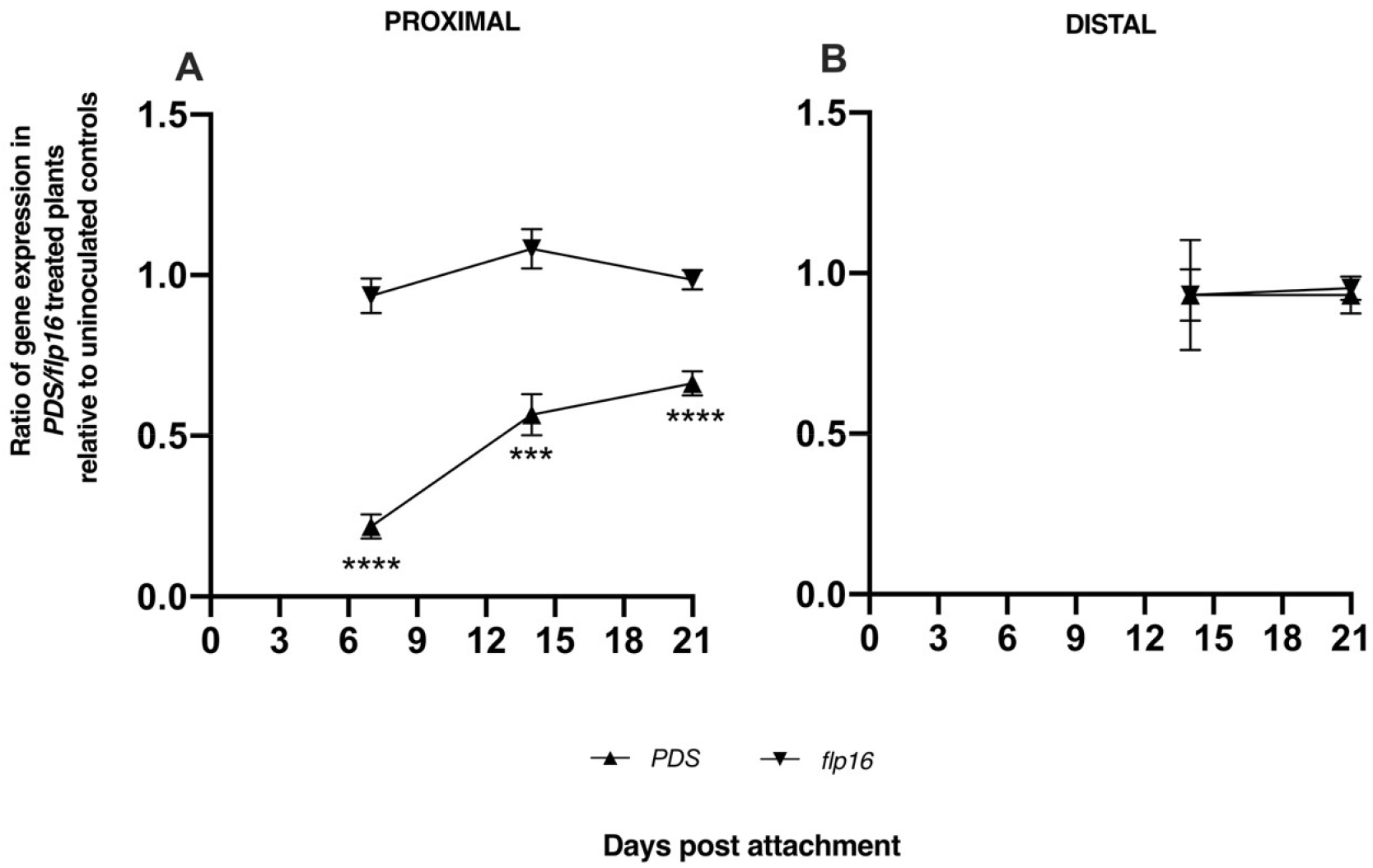
Timecourse analysis of *PHYTOENE DESATURASE* knockdown relative to non-inoculated control plants. Lines represent the ratio of *PDS* transcript abundance in *C. campestris* parasitizing VIGS plants either harbouring pTRV2-*CcPDS* or pTRV2-*Miflp16* at 7, 14 and 21 dpa. (A) shows the relative transcript abundance in the proximal 6cm sections of excised tissue (or whole tissue in the case of day 7). (B) shows the relative transcript abundance in distal 6cm tissues (timepoint 1 is absent as vines at day 7 could not be separated into two distinct sections. Ratios are relative to the endogenous 18s ribosomal gene. Data is representative of 5 biological replicates. An unpaired T-test was used to look for statistical significance between experimental and control plants (*P < 0.05, **P < 0.01, ***P < 0.001).

### Knockdown of *Cc-PDS* reduces *C. campestris* growth

Whole vines were sampled and the lengths and fresh weight were measured. It was consistently observed that there was a significant reductions in total lengths and fresh weight across all time points (Figure 3). The mean total length following PDS knockdown was reduced by 41.62 ± 12.57 mm relative to non-inoculated controls at 7 dpa, 268.1 ± 49.92 mm at 14 dpa and 262.3 ± 62.36 mm at 21 dpa. Similarly, fresh weight was reduced by 11.06 ± 2.85 mg at 7 dpa, 128.1 ± 29.1 mg at 14 dpa and 126.6 ± 41.41mg at 21 dpa.

**Figure 3.**
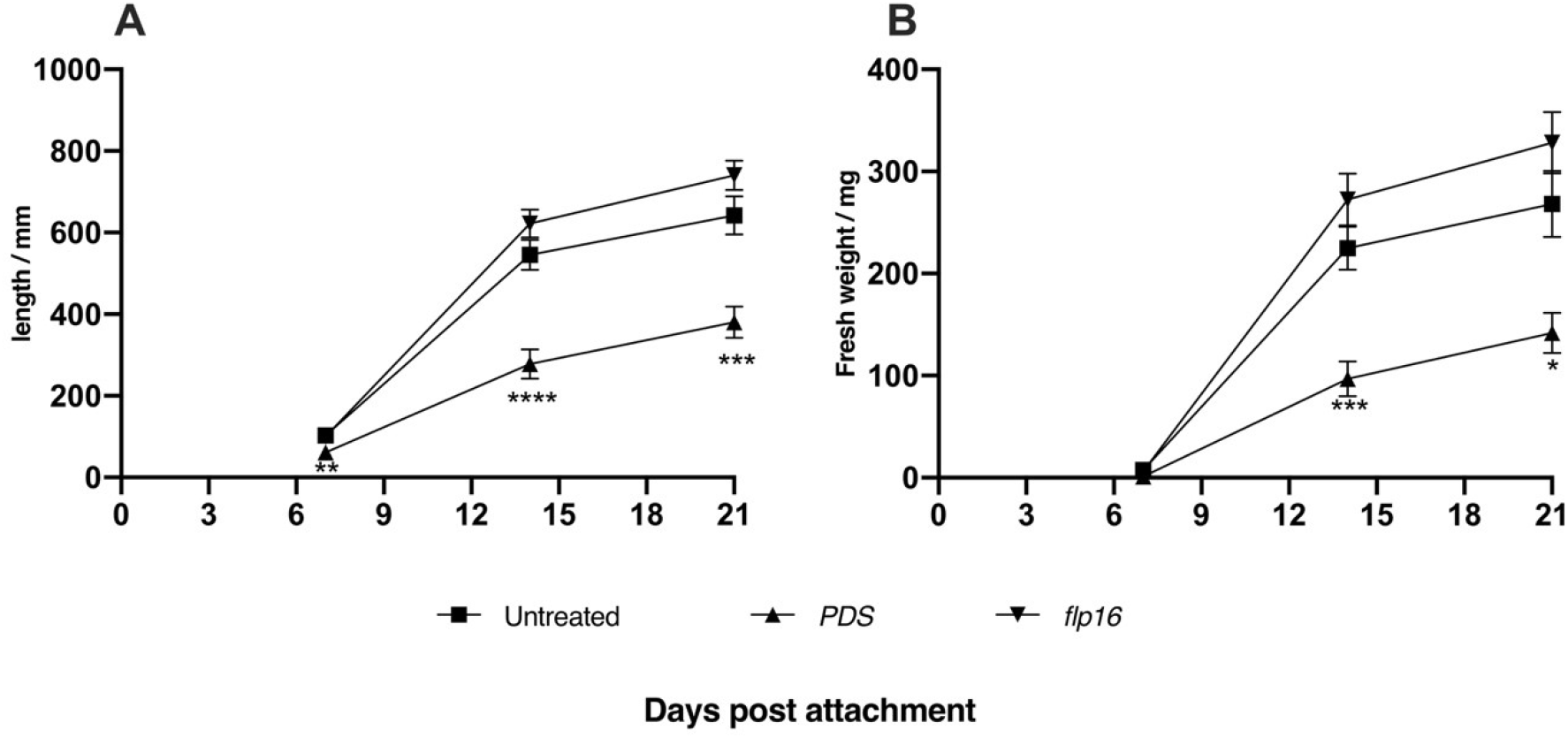
Length and weight of *C. campestris* following infection of VIGS-treated *A. thaliana*. (A) The length of vines excised from host was measured as total length including branched secondary and tertiary vines. (B) The total fresh weight of each individual plant was measured. An unpaired T-test was used to look for statistical significance between experimental and control plants (*P < 0.05, **P < 0.01, ***P < 0.001)

### Silencing of *PDS* in *C. campestris* reduces β-carotene synthesis

Detection of β-carotene content in vines of *C. campestris* was performed using HPLC-MS and quantified relative to a standard curve produced from commercial standards (Sigma). Silencing of *PDS* was concurrent with a significant reduction in β-carotene content in vines at 7 dpa by 8.661 ± 2.229 µg / 100 mg fresh weight and 7.246 ± 2.274 µg / 100mg fresh weight relative to non-inoculation control and flp16 treated samples respectively (Figure 4). At 14 and 21 dpa the levels of β-carotene were still reduced in vines with *PDS* silencing relative to *flp16* and non-inoculated controls but the difference was less severe and not statistically significant. In control vines, tissue β-carotene was observed to decrease as a factor of time.

**Figure 4.**
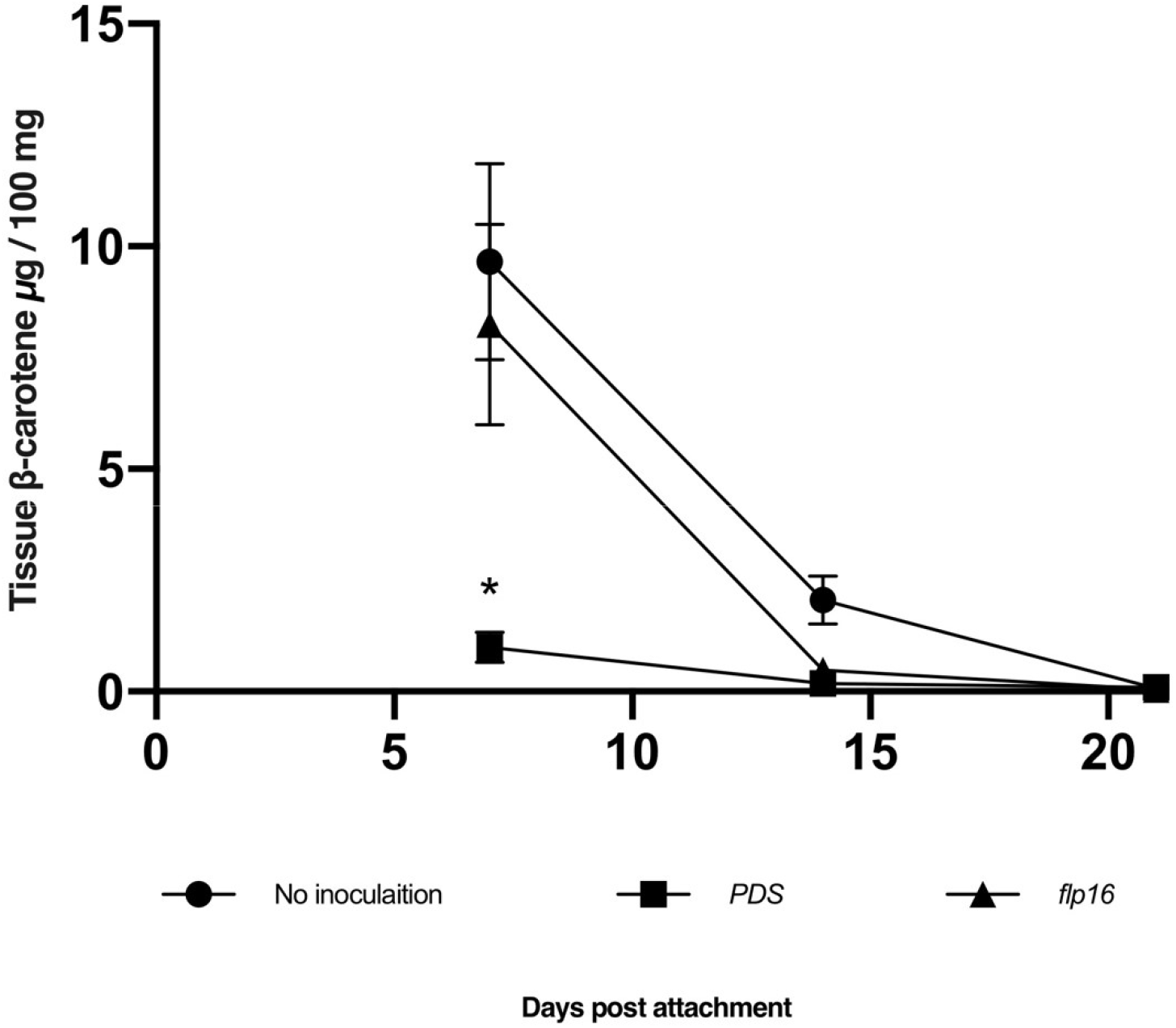
Quantification of fresh weight (FW) β -carotene levels in *C. campestris* over time. Pigment extracts were assayed by HPLC-MS single ion monitoring (API-ES, Pos, frag: 150). Estimates of β -carotene concentration were made by comparison to a standard curve produced from commercial standards and scaled up to the initial sample mass. The mean concentrations at each timepoint were compared by one-way ANOVA and Tukey’s multiple comparisons test (*p<0.05)

## Discussion

The field dodder *C. campestris* is a serious agricultural pest; new approaches to control are urgently required. Whilst it had been demonstrated that *C. pentagona* and *C. campestris* are susceptible to HIGS (Jhu et al., 2021; Alakonya et al., 2012), our data demonstrate for the first time that host-mediated VIGS can effectively reduce the abundance of target genes in *C. campestri*s. The efficacy and ease of VIGS in *C. campestris* paves the way for exploitation of genome sequences by reverse genetics and could rapidly facilitate the development of new herbicidal formulations that rely upon gene-specific dsRNA as the active ingredient.

Whilst we did not attempt to objectively quantify the phenotypic impact of *PDS* knockdown on *C. campestris* branching, we observed an obvious and consistent reduction in branching relative to untreated and VIGS control groups. These observations appear to be consistent with the anticipated impact of *PDS* knockdown on the synthesis of strigolactones (Shinohara et al., 2013).

The applicability of VIGS to the study of *C. campestris* gene function represents a significant advancement in the field and will expedite functional interrogation of the *C. campestris* genome. We anticipate that this will help us to understand host-parasite interactions and could point to new approaches for resistance and control of this economically important parasite.

## Acknowledgements

*C. campestris* ‘doddi’’ seeds were a gift from Prof Michael Axtell (Penn State University).

## Resource availability

The VIGS plasmids used in this study are freely available to other researchers upon request.

